# The mating system of the common house spider *Parasteatoda tepidariorum*

**DOI:** 10.1101/2022.05.27.493702

**Authors:** Apostolos Angelakakis, Natascha Turetzek, Cristina Tuni

## Abstract

Mating systems, with varying female mating rates occurring with the same partner (monogamy) or with multiple mates (polyandry), can have far reaching consequences for population viability and the rate of gene flow. Here, we investigate the mating system of the common house spider *Parasteatoda tepidariorum* (Theridiidae), an emerging model for genetic studies, with yet undescribed reproductive behavior. It is hypothesized that spiders belonging to this family have low re-mating rates. We paired females twice with the same male (monogamy) or with different males (polyandry), scored behaviors and mating success and fitness resulting from single- and double-matings, either monogamous or polyandrous. Despite the study being explorative in nature, we predict successful matings to be more frequent during first encounters, to reduce the risk of remaining unmated. For re-mating to be adaptive, we expect higher fitness of double-mated females, and polyandrous females to experience highest mating success and fitness if reproductive gains are achieved by mating with multiple partners. We show that the majority of the females mated once, not necessarily on their first encounter, and the likelihood of re-mating did not differ between monogamous and polyandrous encounters. The number of matings did not affect fitness, indicated by similar offspring production in females. Female twanging of the web, a behavior that likely advertises female receptivity, lead successful matings, suggesting female control. We discuss how the species ecology, with high mating costs for males and potentially limited female receptivity, may shape a mating system with low mating rates.

## Introduction

Mating systems are primarily shaped by the frequency of mating in each of the sexes, determined by life history and ecological (i.e., socio-demographic) factors (Emlen & Oring, 1977; Andersson, 1994). Female mating rates, whether occurring repeatedly with the same partner (monogamy) or with multiple mates over a single reproductive cycle (polyandry), can have far reaching consequences for population viability and the rate of gene flow (Holman & Kokko, 2013; Pizzari & Wedell, 2013; Lumley *et al*., 2015). Female polyandry appears to be an especially common and widespread mating system (Taylor *et al*., 2014), reported also in socially monogamous species (Westneat & Stewart, 2003). This has challenged the traditional concept of choosy and monogamous females (Bateman, 1948), diverting theoretical and empirical efforts to explain the evolution and maintenance of polyandry (Arnqvist & Nilsson, 2000; Jennions & Petrie, 2000). The main adaptive explanations focus on females receiving direct fitness benefits through increased access to resources (e.g., food, protection) (Arnqvist & Nilsson, 2000) and viable sperm (Reinhardt & Ribou, 2013; Sutter *et al*., 2019), indirect benefits to their offspring by avoiding genetic incompatibility and male genotypes of inferior quality (Jennions & Petrie, 2000; Simmons, 2005), or both (Fedorka & Mousseau, 2002; Tuni *et al*., 2013). Females may also derive direct benefits by accepting multiple matings from reducing the costs associated with male harassment (i.e., convenience polyandry) (Boulton *et al*., 2018). The magnitude of these benefits needs to outweigh the costs associated to mating (e.g., disease transmission, injury, time and energy) (Knell & Webberley, 2004; McNamara *et al*., 2008) for females to solicit re-matings with novel males. Sexual conflict may, on the other hand, represent a non-adaptive route to polyandry where females are forced into a suboptimal number of matings by males (Parker, 2006; Arnqvist & Rowe, 2013).

Despite being ubiquitous, polyandry is highly variable, with extreme numbers of mating partners being reported between and within species (Gowaty, 2013; Taylor *et al*., 2014). For example, honeybee queens *Apis mellifera* mate on average with 12 males during one mating flight (Tarpy *et al*., 2004) and the wasp spider *Argiope bruennichi* with maximum of two (Weiss *et al*., 2020); in a Spanish population of the field cricket *Gryllus bimaculatus* all females were found to be polyandrous (Bretman & Tregenza, 2005), whereas for bumble bees (*Bombus spp*.) 20% of colonies were found to be sired by more than one father (Payne *et al*., 2003). Ecological constraints, such as limited encounter rates between the sexes due to female-biased sex ratios, high mate-search costs for males and/or low population densities, may overall select for low levels of polyandry (Emlen & Oring, 1977; Berger-Tal & Lubin, 2011; Tuni & Berger-tal, 2012). Interestingly, in the absence of paternal care, as in most arthropods, low mating rates are considered to be seldom female-driven (Kokko & Mappes, 2013). Female reluctance in re-mating may instead reflect male manipulation of female behavior (Hosken *et al*., 2009). For example, females of many species enter a refractory period after their first mating that is driven by seminal fluid molecules transferred by males at mating (Chapman & Davies, 2004; Avila *et al*., 2011). The duration of such a period can be substantial (e.g., 30 days for the mosquito *Aedes aegypti* (Degner & Harrington, 2016)), and in short lived animals this may potentially affect lifetime reproduction, leading to single fathered offspring. The fitness benefits uncovered from experimentally inducing polyandry in monogamous females further supports the idea of male-controlled mating rates (Baer & Schmid-Hempel, 1999; Arnqvist & Andres, 2006). If, on the other hand, females maximize their fitness by mating only once, then selection should favor monogamy, which yet, remains an evolutionary puzzle (Klug, 2018).

Most studies testing for adaptive explanations of polyandry estimate fitness benefits of re-mating with same or novel males (Tregenza, 1998; Evans & Magurran, 2000; Simmons, 2005; Slatyer *et al*., 2012). To fully understand how mating systems evolve and are maintained studies should investigate whether re-mating rates are under male or female control by including the study of behaviors at mating for both sexes, together with the fitness consequences of mating decisions. Disentangling whether mate acceptance (or rejections) depends on female choosiness or male manipulation is challenging. Yet studying the outcome of sequential encounters may help shed light on these dynamics. Individuals may behave differently at each encounter, and male-female interactions and the fitness outcomes may largely depend on the outcome of previous encounters (Whittingham & Dunn, 2010). Altogether, these factors may ultimately affect the total number of mating partners in a female’s lifetime.

Spiders represent excellent model organisms to investigate mating system evolution, being well-studied in the field of sexual selection (Eberhard, 2004; Huber, 2005; Elias *et al*., 2011; Tuni *et al*., 2020). Despite most of our knowledge on spider mating rates derives from experimental studies (reviewed in (Tuni *et al*., 2020)), spider mating systems are highly variable, ranging from monogamous wolf spiders (Norton & Uetz, 2005; Jiao *et al*., 2011) to the highly polyandrous nuptial feeding spider (Toft & Albo, 2015). High mating rates may often remain undetected in laboratory studies as females may become sexually reluctant to re-mate after their first mating (Elgar & Bathgate, 1996; Uetz & Norton, 2007; Aisenberg *et al*., 2009), even for extended periods of time (Perampaladas *et al*., 2008). Such decrease in receptivity could indicate increased choosiness in already-mated females or male manipulation of the females’ physiology (Aisenberg & Costa, 2005). The large variation in spiders’ mating systems, alongside being one of the most abundant taxonomical group on earth – with 50.114 reported species (World Spider Catalog, 2022) - calls for more studies comprehensively analyzing species mating behaviors and their fitness outcomes.

In this study we investigate the mating system of the common house spider *Parasteatoda tepidarorium* (previously, *Achaearanea tepidariorum*) of the Theridiidae family (Theridiinae subfamily (Liu *et al*., 2016)). *P. tepidariorum* has become one of the main chelicerate model species used in genetic and developmental biology studies (Hilbrant *et al*., 2012; Oda & Akiyama-Oda, 2020). Benefiting from its small size, relatively short generation cycle and the ability to be raised in large populations under laboratory conditions *P. tepidariorum* is currently one of the best studied spiders in terms of embryonic development and gene family evolution used for comparative evolutionary developmental studies. The molecular experimental methods (including expressional analysis, gene knockdown, overexpression and comparative transgenesis) as well as genomic and transcriptomic data available for this spider are unparalleled by any other chelicerate species (Hilbrant *et al*., 2012; Posnien *et al*., 2014; Turetzek *et al*., 2016; Schwager *et al*., 2017; Oda & Akiyama-Oda, 2020; Janeschik *et al*., 2022). Yet, surprisingly little is known on the species reproductive behavior, although accessible genomic resources and functional gene validation methods would make it an excellent model to understand the genetic basis spider mating behavior. Male-female interactions at mating are only very little studied so far (Ma *et al*., 2022), with records being rather anectodical (i.e., based on observations of only three pairs (Knoflach, 2004)). *P. tepidarorium* builds irregular three-dimensional webs, with a central retreat connected to the surroundings with silk anchor lines (Benjamin & Zschokke, 2003). Males are active and reach sedentary females on their webs, generally staying on the fringe of it, putatively guarding the female from other males or waiting to approach and mate (Ma *et al*., 2022). In this species there is a marked sexual dimorphism, with very small males - male body mass being 10 times smaller than females’ (Oda & Akiyama-Oda, 2020) - that copulate with only few and fast insertions of reproductive organs (i.e., pedipalps) only lasting seconds (Knoflach, 2004). Sexual behaviors have been extensively studied in few species from this Family, with much work done on widow spiders (*Latrodectus spp*.) (Andrade, 1996; Segoli *et al*., 2008; Harari *et al*., 2009; Schraft *et al*., 2021; Sivalinghem & Mason, 2021), and many interesting patterns have emerged, including genital mutilation, in the form of loss of male pedipalps or structures and portions of them (Knoflach & van Harten, 2001) that plug the female genital opening to avoid further inseminations and sexual cannibalism associated to male self-sacrifice behavior (Andrade, 1996; Segoli *et al*., 2008). Records on *P. tepidarorium* document damage of male pedipalps, but mated males are able to nevertheless father offspring, suggesting that the functionality of the pedipalp is not compromised (Locket & Luczak, 1974). While there is lack of visible mating plugs (Knoflach, 2004; Ma *et al*., 2022) the presence of a small apical portion of the male pedipalp was reported inside the female reproductive tract, with its functional role yet unknown (Abalos & Baez, 1963). This often leads to characterizing the species as absent of genital mutilation (Miller, 2007). The rate of sexual cannibalism from laboratory raised females to be relatively low (11-13%) (Ma *et al*., 2022).

It is hypothesized that females belonging to this spider family have low re-mating rates, males being short lived and suffering high mate search costs for sedentary females (Segoli *et al*., 2006), and the timeframe of female receptivity being relatively short (Knoflach, 2004). We therefore tested whether these general predictions also apply to our study species. We paired *P. tepidarorium* females twice with the same male (monogamy) or with different males (polyandry), and scored i) mating behaviors, to describe courtship and mating interactions, ii) mating success, to estimate mating rates, and iii) fitness consequences of single- and double-matings, the latter being either monogamous or polyandrous, to shed light on the adaptive nature of the species mating system. We here provide the first qualitative and quantitative description of mating behaviors that occur during male-female interactions in this species. Although our study is exploratory in nature, we predict successful matings to be more frequent during first encounters with males, to reduce the risk of remaining unmated, and less frequent during subsequent encounters. For re-mating to be adaptive, we expect higher fitness of double-mated females. Specifically, if females gain indirect benefits by mating multiply with different partners, polyandrous matings should be most frequent and yield the highest fitness outcome (i.e., higher hatching success of the brood). If females gain direct benefits (i.e. sperm supply) by mating more than once, we expect no difference in mating success and fitness outcome (i.e., number of offspring) of polyandrous and monogamous matings, but fitness of double-mated females to be higher than for single mated females.

## Materials and Methods

### Collecting and rearing spiders

Spiders were either captured from several buildings located in Munich (Germany) during March and April 2021 or taken from laboratory stock lines derived from the original genome lines (Schwager *et al*., 2017) (received from Göttingen, (Janeschik *et al*., 2022)). This allowed reaching an adequate sample size of 183 spiders, 125 originating from the genome line (hereafter, G) and 58 from wild-caught females (hereafter, W), and variation in origin was accounted statistically. All spiders were raised from egg emergence to adulthood in the laboratory at room temperature (approx. 23 C°) under natural light conditions. Cocoons laid by females, were collected two days after egg-laying and placed in a new vial covered with a styrofoam plug and equipped with tissue paper on the bottom that was moisturized three times a week. After hatching spiderlings were fed with approximately 20-25 fruit flies (*Drosophila sp*.). Ten days later, spiderlings were separated into individual vials (2.5 x 9.5 cm) and were fed five fruit flies, three times per week. Once juveniles (2-3^rd^ instar), 10 fruit flies were provided with the same frequency. Females that reached a subadult stage (i.e., 5^th^ instar) were transferred to bigger vials (5 x 10 cm) and given 1-2 house flies (*Musca domestica*), while males were instead given 7-8 fruit flies three times per week throughout their adulthood. Sexes of this species can only be distinguished once they undergo several (5-6) molting stages (Quade *et al*., 2019), and become sexually dimorphic. Females retain a larger abdomen (i.e., opisthosoma) and remain stationary on webs, while males, which are smaller in size, possess a slimmer, red-colored body, develop a pair of thick pedipalps and display active walking behavior.

### Mating trials

Spiders were randomly assigned to mating trials 1-8 weeks post maturation to adulthood. There was no significant difference in age at testing (N days from adulthood to 1^st^ trial; mean N days (g) ± SE) of females (monogamy 17.35 ± 0.92, n=40; polyandry 17.22 ± 1.03, n=41; t-test, log-transformed, t=-0.76, df=1, n=81, p=0.44), but males assigned to the polyandrous treatment were unintentionally younger than those in the monogamous (monogamy 26.26 ± 2.14, n=38; polyandry 16.86 ± 1.81, n=29; t-test, log-transformed, t=-3.47, df=1, n=67, p=0.001*), which we controlled for statistically (see below). Before testing, male vials were inspected for the presence of a sperm web, which generally appeared as a horizontal web sheet on the styrofoam plug of the vial. Male copulatory organs in spiders are not connected to the testes, and through the process of sperm induction, males release sperm on sperm webs and uptake them in their pedipalps by dipping these in the sperm droplet. This process is known to last for relatively long in *P. tepidarorium* (Knoflach, 2004) and is not repeated during copulation. It was observed only occasionally, hence the presence of the web was used as a proxy to consider sperm induction completed in males.

In order to limit stressful manipulation and damage of female webs we measured individual body mass to the nearest 0.001 mg two days before the first mating trial using a semi-micro digital scale (Mettler Toledo). Females were exposed sequentially to two males, once per day with a one-day break: they were either given the same male twice (monogamy treatment, n= 40 females) or two different males (polyandry treatment, n = 44 females). The males matched the female’s social experience (i.e., a naïve male and female were paired in trial 1, an experienced male and female on trial 2), with those from the polyandrous treatment being used with two different females. In the case of sexual cannibalism, where the female killed and consumed the male, the latter was replaced with a novel male. Therefore, males were tested 1-2 times each, with the exception of one male used 3 times. Individuals assigned to a mating pair were never from the same cocoon to avoid inbreeding (i.e., crossing siblings). Animals derived from G were 60 females and 65 males, and those from W 24 females and 34 males. Pairs consisted of G females paired twice with G (39.3%, n= 33) or W (25%, N=21) males, and W females paired twice with G (22%, n=19) and W (8%, n= 7) males, while in 4 cases these mated with both W and G (6.7%).

Trials were conducted by placing the male inside the female’s housing vial, by lifting the lid of the vial on one side and gently pushing the male inside with a paintbrush. The vial was then located inside a self-made chamber with carton side walls and top to limit environmental disturbance. A light-source was placed stationary on the back side to allow video recordings of male and female interactions. A GoPro Hero9-Camera was placed in front of the vial at a fixed distance of 20 cm. Spiders were recorded for a total of 20 minutes. Occurrence of cannibalism and mating were noted by the observer (A.A.). After the trial, males were returned to their housing vial.

### Fitness

Once females completed the two mating trials, regardless of their outcome they were monitored daily for cocoon production for 4 consecutive weeks. Females of this species lay up to 10-12 cocoons, but due to our methodological approach they produced a maximum of 4 cocoons.

When laid, cocoons were collected in standard glass test tubes and the wrapped silk was opened after 5 days using Dumont #5 pair of forceps (Fine Science Tools). Eggs were kept in test tubes with a small piece of wet paper on the lid. These were checked daily if they have reached the postembryonic stage (approximately 180 hours after laying of the cocoon at 25°C (Mittmann & Wolff, 2012)). After hatching the total number of nymphs, unhatched eggs, as well as the “unsuccessful” eggs, - shrunken smaller eggs compared to the others eggs with a light-yellow color - were counted using a stereomicroscope (Zeiss). This allowed estimating the proportion of hatched eggs per cocoon.

### Video scoring of mating behaviors

We scored videos of the mating trials to qualitatively and quantitively describe male and female mating behaviors using the software BORIS. During male-female interactions, males generally approach the female on her web by walking towards the female, and performing tapping movements with front legs, seemingly in response to certain female leg movements and vibrations on the web. Specifically, the female repeatedly bounces her abdomen and, synchronously or not, lifts legs I and II to pluck web strings, a behavior defined as twanging in other Theridiids (*Parasteatoda wu* (Lubin, 1986)), or simply plucking (Knoflach, 2004). Males approach the female and often interact physically by touching the female with their anterior legs. They may also respond with vibrations (i.e., bouncing of abdomen and/or body rocking) themselves. These behaviors are repeated in multiple sequences until successful pairing occurs. Copulation is fast, after positioning below the female the male inserts his pedipalp into the female epigyne and within seconds releases his sperm by applying hemolymph pressure indicated by swelling of the pedipalps tip (Knoflach, 2004). After sperm transfer the male directly moves away from the female.

For each video we scored the following behaviors: 1. time from the start of the trial until first female twanging; 2. total number of female twanging; 3. time until first male approach, by moving towards the female and making physical contact with front legs; 4. total number of male approaches; 5. total number of attempted matings (i.e., male tries to enter the mating position); 6. occurrence of female cannibalism (i.e., female consumption of the male, pre or post-mating); 7. mating (i.e., male enters mating position and inserts pedipalp), and whether it occurs once or twice during the same trial; and 8. latency to mate (i.e., time from the start of the trial to successful mating). We did not score bouncing due to the difficulty in scoring such behavior accurately from the videos. Among the total 168 videos, for 31 video-inspection was not possible (e.g., damaged or missing video-recordings). In 6 trials females failed to build a functional web, instead loosely spinning few silk threads on the bottom of the vial. Therefore, behaviors could not be scored as males failed to approach the female without a web and the female failed to signal to the male. The outcome of matings in the latter was always unsuccessful and the datapoints excluded from further analyses.

### Statistical analyses

To test whether mating success (i.e. successful copulation of the pair), male, and female behaviors at mating are affected by the treatment applied, we conducted generalized linear mixed effect models (GLMMs) including as fixed effects treatment (monogamy and polyandry), trial order (first and second), their interaction, and relative body mass difference between the sexes (female mass - male mass), which could play a role in male-female interactions. Male and female ID were included as random effects. Binomial family distribution was used for mating success, Poisson for latencies. The relative body mass difference and the trial order were grand-mean-centered. To specifically ask if re-mating rates with same (monogamy) or different (polyandry) males, were affected by the spiders’ previous mating experience (i.e., outcome of trial 1) we ran a linear model testing for the effect of a previous mating (successful or not), treatment (monogamy and polyandry), and body mass difference on the likelihood of successful matings during trial 2. To account for variation in spider origin (wild-caught, W and laboratory genome line, G) and in male age, which due to logistics was not standardized, we expanded the statistical models described above to include these as additional factors.

We tested the correlation between number of female twangings and male approaches using correlation coefficient, as these appeared to be consequential. Finally, we tested for the effect female twanging on the outcome of the mating (successful copulation or not) by re-running the GLMM for analysing mating success described above, including female twanging. Female twanging was grand-mean-centered and normalized with the standard-deviation. Due to the paucity of data occurrence of cannibalism in monogamous and polyandrous and first and second trials, was analysed using Chi-square test. T-tests were used to test for differences in relative body size difference and female twanging behaviour in trials with and without cannibalism. We also tested if the number of attempted matings differ between mated and unmated pairs using t-test.

To test whether mating once or twice, with the same (twice monogamy, M) or different male (twice polyandry, P), affected female fitness we conducted linear models to analyze the effect of treatment (mated once, twice M, twice P) and female body mass on the total number of cocoons laid (0-4), the proportion of viable cocoons (n viable cocoons/total N cocoons) and the total number of offspring produced per female. Generalized linear mixed effect models were used to further explore the effects of treatment (mated once, twice M, twice P), female mass and cocoon number (1-4) on the number of eggs laid per cocoon and the proportion of eggs that hatched per cocoon. Female ID was included as a random effect. The model was further expanded to include spider origin (see above).

## Results

### Mating rates

We observed overall low mating rates, as a total of 39 (45.2%) females and 51 (51.5%) males mated successfully (61 trials out of 168). The number of females that mated once was significantly higher than those mated twice (χ^2^ = 4.33, df = 1, p<0.037; **Figure 1**). No significant difference in mating success was observed between females mating twice with the same (twice monogamy, 15% n=6) or novel mate (twice polyandry, 16%, n=7) or (χ^2^ =0.077, df=1, p=0.78; **Figure 1**). The likelihood of mating successfully during a trial was not affected by mating treatment (31.5%, n=24, monogamy; 44%, n=38, polyandry), nor by the order of testing (46.2%, n=37 in trial 1; 30.5 %, n=25 in trial 2) (**Table 1; Figure 2**), or the relative difference in body mass between the sexes (**Table 1**).

**Table 1.** Results from the GLMMs investigating the effects of mating treatment (polyandry and monogamy), mating trial number (1 and 2), their interaction, and relative body mass difference between the sexes (female-male mass) **(a)** and intensity of female twanging behaviour (N times performed) **(b)**, on the likelihood of mating successfully (GLMM binomial). Point estimates and 95% credible intervals are shown on a logit scale and relative to the reference category (Intercept, monogamy, and for the other effects in a standardized level). The residual variance component is fixed to π^2^/3. Significant effects are shown in bold.

|  | **Mating success** |  |
| --- | --- | --- |
|  | **Model a)** | **Model b)** |
| **Fixed effects** | **$\beta$ (95% CI)** |  |
| (Intercept)* | -0.87<br>(-1.42, -0.33) | -1.05<br>(-1.68, -0.47) |
| Treatment (polyandry) | 0.61<br>(-0.18, 1.390) | 0.47<br>(-0.44, 1.41) |
| Mating trial (1,2)<sup>a</sup> | -0.54<br>(-1.56, 0.47) | -0.29<br>(-1.39, 0.73) |
| Treatment (polyandry) X Mating trial<sup>a</sup> | -0.41<br>(-1.79, 0.92) | -1.23<br>(-2.86, 0.41) |
| Relative Body Mass<sup>a</sup> | -0.08<br>(-13.27, 13.77) | -0.70<br>(-16.95, 15.31) |
| N female twanging behavior <sup>b</sup> |  | **0.69**<br>**(0.23, 1.18)** |
| **Random Effects** | **$\sigma^2$(CI 95%)** |  |
| Female ID | 0.50<br>(0.37, 0.67) | 0<br>(0 , 0) |
| Male ID | 0<br>(0 , 0) | 0.32<br>(0.23, 0.42) |

\* Reference category, estimate for treatment monogamy and mean values of remaining fixed effects
<sup>a</sup> mean centered
<sup>b</sup>mean-centered and normalized with the standard-deviation

**Figure 1.**
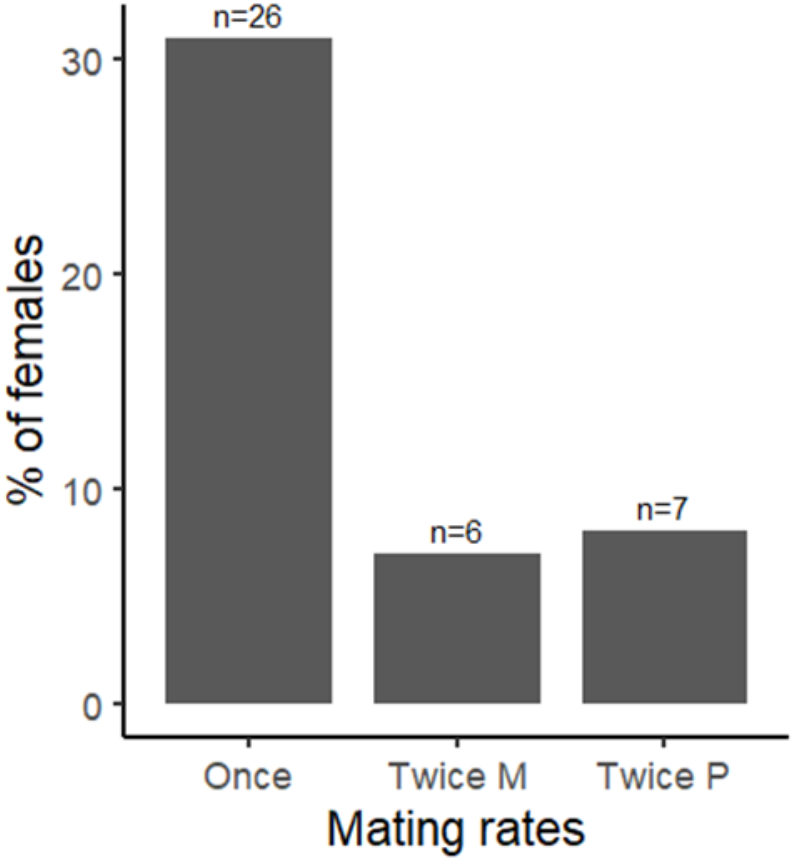
Females (%) that successfully mated only once, twice with the same mating partner (Twice M = monogamy) and twice with a novel mating partner (Twice P=polyandry).

**Figure 2.**
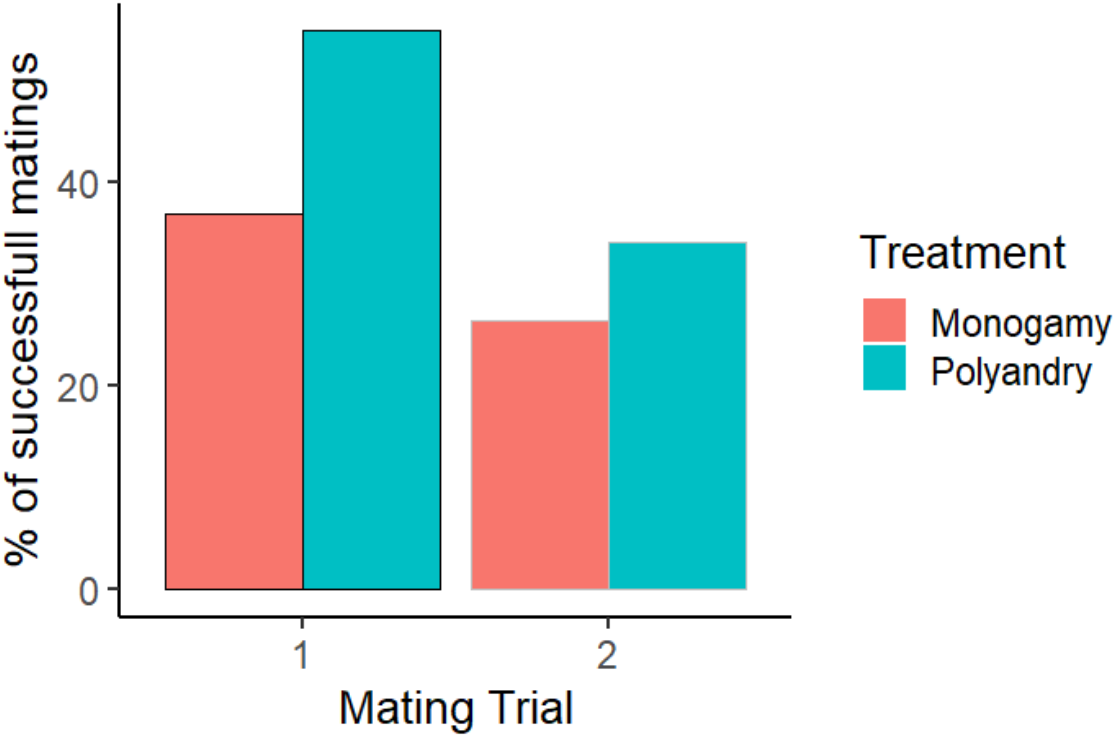
Successful matings (%) during 1^st^ and 2^nd^ trials, for spiders in monogamous and polyandrous treatments

The likelihood of re-mating (i.e., trial 2) was not affected by the novelty of the mating partner, whether the same male (monogamy) or a novel male (polyandry) or by the outcome of the previous mating trial, whether successful or unsuccessful mating (GLM - binomial, β (95% CI); Intercept* −1.23 (−2.09, −0.36); Treatment 0.21 (−0.82, 1.30); Previously mated (yes, no) 0.59 (−0.39, 1.55); Relative body mass −2.74 (−20.42, 15.33)). Male age and spider origin (W or G) did not affect likelihood of mating nor re-mating (Appendix **Table S1**).

### Behaviors at mating

Pre-copulatory cannibalism occurred in 17 trials (10.4%), with the only exception of one occasion of post-copulatory cannibalism (trial 1, polyandry treatment) and did not differ between mating treatments (n=4, monogamy; n=13, polyandry; χ^2^=3.18, df=1, p=0.07), testing order (n=10, trial 1; n=7, trial 2; χ^2^=0.32, df=1, p=0.57). The relative difference in body mass between the sexes was higher in non cannibalised pairs compared to cannibalised pairs (Welch 2 sample t-test, t= 2.32, df= 26.48; p = 0.028).

None of the behaviours measured – time to first female twanging, total number of female twangings, time to first male approach, total number of male approaches, time to mating - were affected by the mating treatment (monogamy and polyandry), nor body mass difference between the sexes (**Table 2 and 3)**. In contrast an interaction between treatment and testing order (trial 1 and 2) affected female behaviors as those of the polyandrous treatment performed twanging later in time and performed less twanging during their 2^nd^ trials. Similarly, males of the polyandrous treatment approached females later in time and less frequently on their 2^nd^ trial (Table 2). Male age and spider origin (W or G) did not affect any of the above-mentioned behaviors (Appendix **Table S2**).

**Table 2.** Results from the GLMMs investigating the effects of mating treatment (polyandry and monogamy), mating trial (1 and 2), their interaction, and relative body mass difference (female-male mass) on male and female behaviours. Point estimates and 95% credible intervals are shown on a logit scale and relative to the reference category (Intercept, monogamy, and for the other effects in a standardized level). The residual variance component is fixed to π^2^/3. Significant effects are shown in bold.

|  | Time to female twanging | Total N of female twanging | Time to 1<sup>st</sup> male approach | Total N male approaches | Time to mating |
| --- | --- | --- | --- | --- | --- |
| **Fixed Effects** | **$\beta$ (95% CI)** |  |  |  |  |
| (Intercept)* | 4.34<br>(3.89,4.76) | 2.90<br>(2.50,3.29) | 5.03<br>(4.59,5.49) | 1.45<br>(1.03,1.86) | 506.56<br>(342.47,664.19) |
| Treatment (polyandry) | 0.16<br>(-0.47,0.85) | -0.36<br>(-0.91,0.22) | -0.19<br>(-0.89,0.48) | -0.19<br>(-0.78,0.42) | -26.93<br>(-306.75,239.10) |
| Mating trial (1,2)<sup>a</sup> | 0.04<br>(-0.01,0.08) | -0.06<br>(-0.14 , 0.04) | **0.08**<br>**(0.05 , 0.12)** | **-0.25**<br>**(-0.43 , -0.05)** | 61.00<br>(-237.93,354.29) |
| Treatment (polyandry) x Mating trial <sup>a</sup> | **0.78**<br>**(0.65,0.89)** | **-0.41**<br>**(-0.63,-0.19)** | **0.87**<br>**(0.73,1.03)** | **-0.45**<br>**(-0.84,-0.04)** | -289.19<br>(-776.26,194.06) |
| Relative Body Mass<sup>a</sup> | -7.03<br>(-17.67,4.28) | 8.01<br>(-1.82,17.97) | 0.96<br>(-10.02,12.37) | 4.16<br>(-6.15,14.77) | -3155.99<br>(-8319.68,1682.37) |
| **Random Effects** | **$\sigma^2$(CI 95%)** |  |  |  |  |
| **Female ID** | 0.72<br>(0.52,0.96) | 0.32<br>(0.21,0.44) | 0.45<br>(0.31,0.64) | 0.24<br>(0.16,0.33) | 0<br>(0,0) |
| **Male ID** | 1.18<br>(0.92,1.54) | 1.24<br>(0.95,1.59) | 1.43<br>(1.07,1.89) | 1.39<br>(1.06,1.78) | 17706.46<br>(8393.57, 31639.56) |

\* Reference category, estimate for treatment monogamy and mean values of remaining fixed effects
<sup>a</sup> mean centered

**Table 3.** Estimates for each behaviour given in mean ± SE, median, range and sample size (N).

|  | **Monogamy** |  | **Polyandry** |  |
| --- | --- | --- | --- | --- |
| **Behaviours** | **Trial 1** | **Trial 2** | **Trial 1** | **Trial 2** |
| Time to 1<sup>st</sup> female twanging (secs) | 137.24 ± 41.02,<br>50.29,<br>2.41-1064.53,<br>32 | 166.48 ± 46.43,<br>50.9,<br>5.91- 1200.62,<br>35 | 150.48 ± 48.76,<br>64.94,<br>11.90-1063.08,<br>25 | 280.84 ± 71.58,<br>73.4,<br>5.15-1065.73,<br>25 |
| Total N of female twanging | 28.5 ± 3.96,<br>26,<br>0-106,<br>36 | 25.95 ± 4.29,<br>13,<br>0-98,<br>37 | 22.41 ± 4.41,<br>15,<br>0-107,<br>29 | 22.07 ± 3.67,<br>22,<br>0-61,<br>27 |
| Time to 1<sup>st</sup> male approach | 249.35 ± 58.85,<br>122.77,<br>7.41-1163.02,<br>25 | 279.71 ± 52.86,<br>208.91,<br>8.90-1173.34,<br>28 | 161.84 ± 55.56,<br>81.90,<br>4.41-1093.91,<br>20 | 284.21 ± 75.15,<br>97.91,<br>19.91-1088.74,<br>21 |
| Total N male approaches | 7.39 ± 1.12,<br>8,<br>0-23,<br>36 | 6.08 ± 1.14,<br>4,<br>0-25,<br>36 | 7.86 ± 1.60,<br>7,<br>0-34,<br>29 | 6.11 ± 1.21,<br>4,<br>0-21,<br>26 |
| Time to mating (secs) | 452.15 ± 56.31,<br>503.17,<br>33.43- 1127.02,<br>12 | 524.95 ± 50.39,<br>452.32,<br>202.61- 999.98,<br>10 | 617.28 ± 53.74,<br>581.30,<br>82.45- 1156.74,<br>12 | 412.41 ± 60.94,<br>286.18,<br>79.65-997.65,<br>4 |

Females performed twanging in 89% of the trials (117 out of 131 video-scored trials), and males approached females in 76% of the trials (100 out of 131). Males approached females only after females performed twanging (mean time to male first approach for males 248.58 ± 30.10 seconds, n=100; mean time to first female twanging, 179.50 ± 25.86 seconds, n=117). The two variables are positively correlated; the sooner the female started twanging the sooner male approached (R= 0.53, p= <.0001, n=97; **Figure 3a**) and the total number of times female twanging was performed correlates positively with the number of male approaches (R=0.61, p= <.0001, n=96, **Figure 3b)**. Higher numbers of female twanging significantly affected mating success (**Table 1; Figure 4a**). Females performed significantly more twanging during trials in which males were not cannibalized (t-test, t=6.83, df=42.1, p<0.01; Figure 4b).

**Figure 3.**
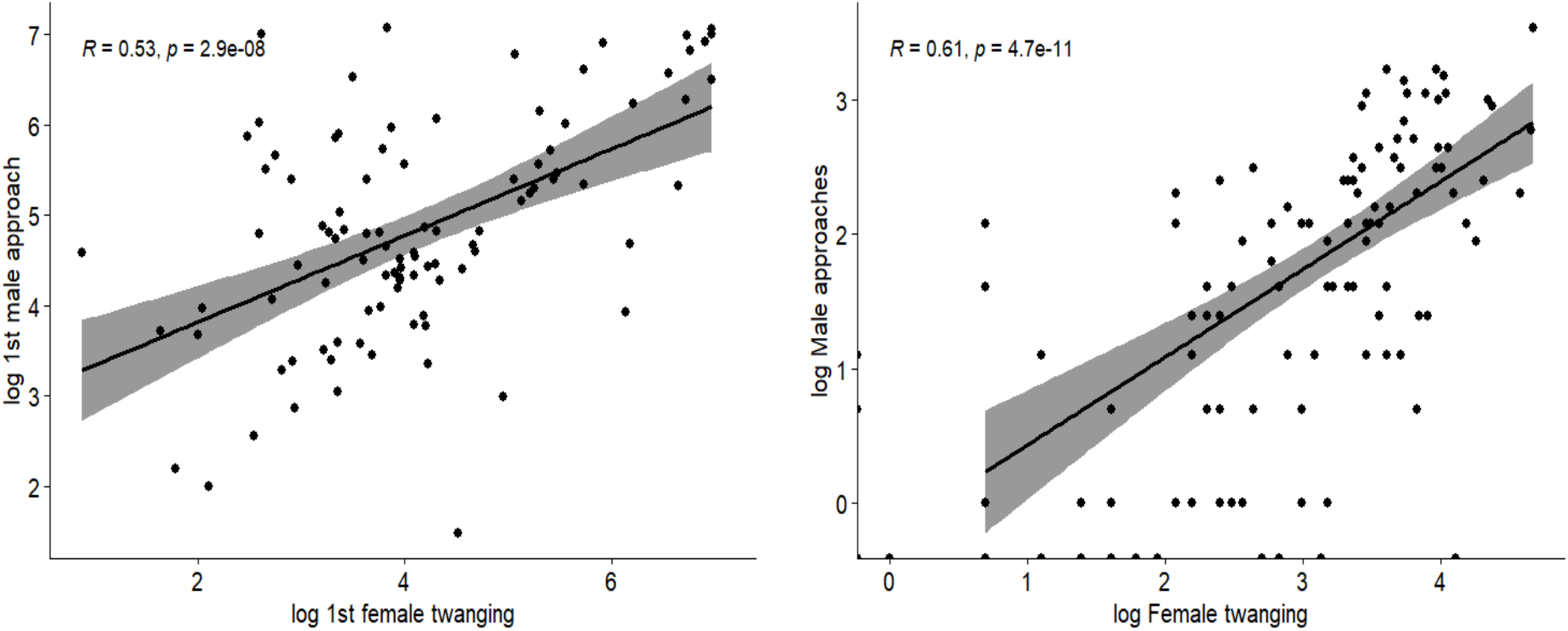
**a)** Positive correlation between the time until first female twanging and first male approach, and **b)** between the total number of female twanging and number of male approaches.

**Figure 4.**
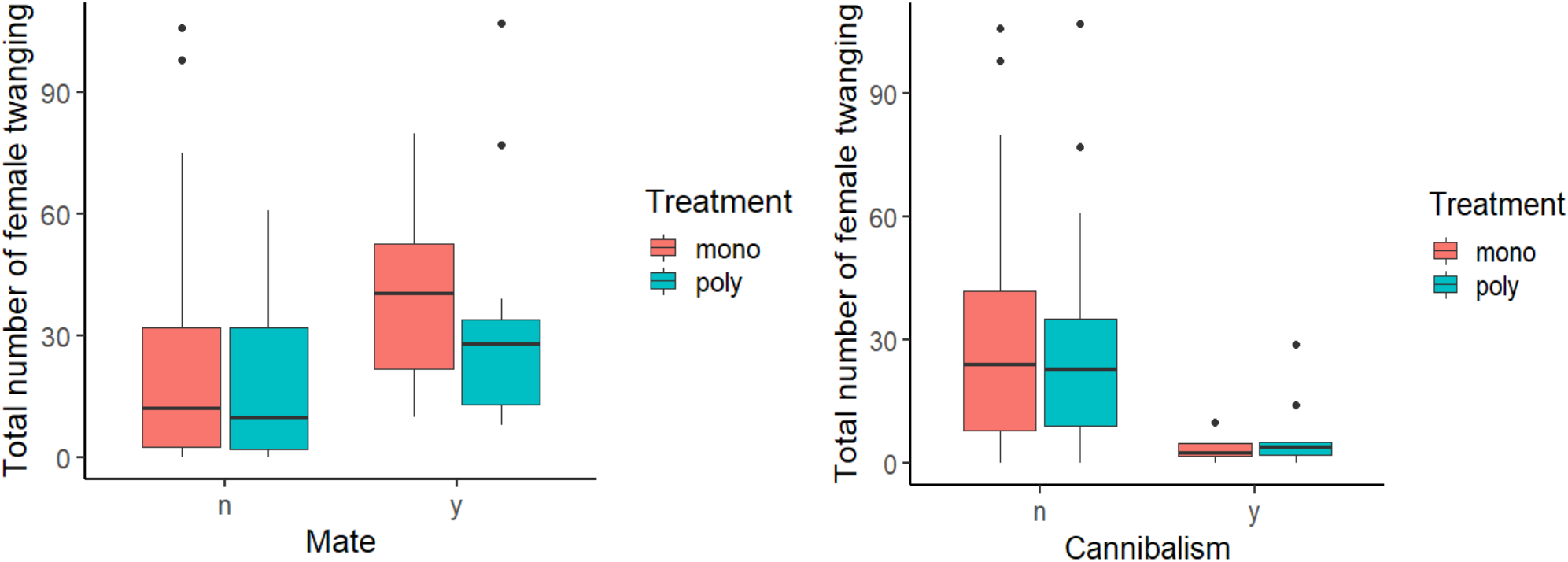
In trials where successful matings occurred (a) and in trials where males were not cannibalized (b), females performed a higher number of twanging.

The number of mating attempts are higher in the trials with successful matings (8.30 ± 1.1, n=43) compared to unsuccessful (3.95 ± 0.79, n=84) ones (t-test, t= 3.24, df=81.28, p=0.0017). In 8 trials males entered the mating position twice, meaning they re-approached the female a second time and were able to couple their pedipalps again during the 20-minute trial. These were distributed as follows: 4 trials from the monogamous treatment (three during the trial 1 and one during trial2) and 4 trials from the polyandry treatment (two during trial 1 and two during trial 2).

### Fitness

One female from the polyandrous treatment died 2 days after the trials, leaving us with a sample of 83 females. Successful reproduction (where at least one cocoon hatched) occurred in 88.23% (n=30) of females mated once, 66.66% (n=4) of those mated twice monogamously and 87.5% (n=7) of those mated twice polyandrously. The total number of cocoons laid by mated females (0-4) within 4 weeks, the proportion of those that hatched as well as the total number of viable offspring, were not significantly affected by the mating treatment (once, twice M, twice P), or female body mass (**Table 4 and 5**).

**Table 4.** Estimates for fitness measures given in mean ± SE, median, range and sample size (N).

|  | **Mated once** | **Mated twice monogamy** | **Mated twice polyandry** |
| --- | --- | --- | --- |
| **Total number of cocoons** | 3.42± 0.23,<br>4, 0-4,<br>26 | 3 ± 0.63,<br>3.5, 0-4,<br>6 | 3.57 ± 0.3,<br>4, 2-4,<br>7 |
| **Proportion of viable cocoons laid** | 0.82± 0.06,<br>1, 0-1,<br>26 | 0.68 ± 0.17,<br>0.75, 0-1,<br>6 | 0.77 ± 0.14,<br>1, 0-1,<br>7 |
| **Total number of offspring** | 353.38± 44.09,<br>376, 0-819,<br>26 | 224.17± 77.95,<br>263, 0-421,<br>6 | 361.57± 114.22,<br>314, 0-921<br>7 |

**Table 5.** Results from the GLMs investigating the effects of mating treatment and female body mass on fitness.

| **Model Estimates $\beta$ (95% CI)** |  |  |  |
| --- | --- | --- | --- |
|  | **Total number of cocoons** | **Proportion of viable cocoons laid** | **Total number of offspring** |
| **(Intercept)\*** | 4.50<br>(3.38, 5.66) | 0.81<br>(0.47, 1.16) | 383.62<br>(144.58, 630.38) |
| **Mated twice monogamy** | -0.32<br>(-1.31, 0.72) | -0.14<br>(-0.46, 0.20) | -125.20<br>(-336.05, 95.13) |
| **Mated twice polyandry** | -0.12<br>(-1.09, 0.85) | -0.05<br>(-0.34, 0.24) | 1.49<br>(-205.27, 206.47) |
| **Female Body Mass** | **-13.66**<br>**(-26.98, -0.09)** | 0.16<br>(-3.98, 4.38) | -392.34<br>(-3224.81, 2494.56) |

The number of eggs laid per cocoon and the ratio of hatched eggs (the number of hatched eggs/the total number of eggs per cocoon) per cocoon were not affected by the mating treatment (mated once, twice M, twice P), laying order of the cocoon (n of cocoon) and female mass (Appendix **Table S3)**.

## Discussion

Our study was designed to gain insights on the behavior and mating system of the emerging spider model system *Parasteatoda tepidariorum*. We revealed that the majority of the females only mate once, and not necessarily on their first encounter (Figure 1). In addition, for the few females that mated twice there was no preference for re-mating with the same male (monogamy) or with a novel male (polyandry) (Figure 2). Beyond that, we observed that mating once or twice, regardless of whether monogamous or polyandrous, did not affect female fitness, indicated by similar offspring production in all tested females. In summary our findings suggest that *P. tepidariorum* possesses a mating system characterized by low mating rates.

### Low mating rates in Parasteatoda tepidariorum

The low mating rates reported are striking. On the one hand, these may be a consequence of high choosiness of females. Mate choice may provide females with reproductive benefits (Rosenthal & Rosenthal, 2017), despite mate sampling entails energetic costs and the risk of reproductive failure (Kokko & Mappes, 2005). In this respect, one of the general expectations derived from sexual selection theory is, that females are less choosy when unmated, accepting copulations from the first male they encounter to avoid the cost of remaining unfertilized (Andersson & Simmons, 2006; Bleu *et al*., 2012; Tanner *et al*., 2019). On the other hand, selection on males to reduce sperm competition may likely be responsible for the evolution of male manipulative adaptations, such as seminal substances that can act on female reproductive behavior and physiology to lower female receptivity (Chapman & Davies, 2004; Aisenberg & Costa, 2005). Interestingly the likelihood of mating successfully in *P. tepidariorum* females did not differ during first or subsequent encounters with males, nor was affected by the female’s mating status (unmated or previously mated). We can therefore exclude unreceptivity and/or enhanced choosiness of once-mated females as mechanistic explanations for our findings. We, however, cannot entirely rule out that mate choice for traits other than those accounted for in our study may have driven the large numbers of unsuccessful matings. Courtship vibrations in web-building spiders are known to convey information on the signaler and its underlying quality (Herberstein *et al*., 2014), lowering female aggressiveness and influencing female mate choice (Wignall & Herberstein, 2013a; b). For example, vibrations performed on cobwebs from widow spiders are suggested to convey information on male size (Sivalinghem & Mason, 2021). We were unable to reliable quantify vibratory movements in the form of male bouncing of the abdomen and/or body rocking, which would have nevertheless informed very poorly on the transmission of vibratory signals to females.

The lack of successful matings also do not appear to be caused by non-receptive individuals. Matings occurred in 36% of the trials. However, females signaled in 89% of the trials and males actively approached females in 76% of the trials. Based on our personal observations, together with reports from (Ma *et al*., 2022), we also note that females occasionally cooperate with males by presenting the opening of their genitalia tract to the courting individual. Yet, the number of mating attempts required before males could successfully enter the mating position and transfer sperm, within a 20-minute trial, was high (on average 8). Despite an extensive overview of 70 species of Theridiids reported that courtship lasts on average 24 minute in comb footed spiders belonging to the Steatoda-type, which includes *P. tepidariorum* (Knoflach, 2004), we suggest that males in this species may require longer to interact with the female. These mating attempts may serve as male courtship and exchange of information between the sexes.

Ecological constraints that limit the encounter rates between the sexes, such as high mortality and mate-search costs for males (Byers *et al*., 2006; Kasumovic *et al*., 2006), low population densities (Xue *et al*., 2016), or the spatial and temporal distribution of receptive females (e.g., female-biased sex ratios) (Emlen & Oring, 1977; Ims, 1988; Tuni & Berger-tal, 2012), may also all contribute to shaping mating systems with low levels of polyandry (Elias *et al*., 2011). To our knowledge, data on population structure (e.g., sex ratios) and natural encounter rates between males and females are missing for this species. We know that mate-search is risky in spiders (Kasumovic *et al*., 2006; Berger-Tal & Lubin, 2011). Male theridids, in particular, develop into adults and leave their webs in search of sedentary females facing high mortality as reported in several widow spiders (Andrade, 2003; Segev *et al*., 2003; Segoli *et al*., 2006; Scott *et al*., 2019). An indication of such costs is for example reported in males of the western black widow *Latrodectus hesperus* that, with an 88% mortality during mate search, parasitize the mate-search effort of other males in the population to more efficiently find females (Scott *et al*., 2019).

Generally once spider females mature, they signal their receptivity using sex pheromones, emitted from their body or the silk of their webs (Kasumovic & Andrade, 2004; MacLeod & Andrade, 2014). The timeframe of female receptivity is generally short and this imposes further constraints on mating rates, considering also that males generally become sexually mature (i.e., molt to the adult stage) before females (Elias *et al*., 2011). Low mating rates may also result from monygy, where males mate with only one female (Schneider & Fromhage, 2010). Such reproductive strategy is facilitated by sexual cannibalism (e.g., self-sacrifice) and genital mutilation, because of strong sexual selection on competing males (Fromhage *et al*., 2005), and has been shown to occur in theridids (Michalik *et al*., 2010). The risk of pre-copulatory sexual cannibalism, performed in 10% of the trials from well-fed females in our study, may further act on constraining male mating attempts. Finally, *P. tepidariorum* lacks forms of male adaptations allowing for female monopolization and limiting inseminations from additional males, such as extremely long copulations that function as mate guarding (Simmons, 2001), or visible plugging of the outer female genitalia (Uhl *et al*., 2010) which would represent adaptations to high male-male competition. In summary, all the above-mentioned factors may select for low polyandry in this study species.

#### Re-matings do not lead to increased fitness

Low mating rates and monogamy may be favored when increased mating rates have little effect on female reproductive output (Klug, 2018). We could clearly show that re-matings did not provide females with either direct or indirect fitness benefits. Indeed, females mated once had similar cocoon and offspring production success as double-mated females lasting for up to 4 weeks post-mating. This indicates that females of this species are able to maximize their fitness by mating only once. Despite low sample sizes warrant cautious interpretation, we also found no differences in the reproductive output of females mated monogamously and polyandrously. The benefits of polyandry in spiders have been detected in the form of increased female fecundity (Uhl *et al*., 2005) or resource acquisition (i.e., foraging advantages) (Watson, 1993), but also as benefits to their offspring (Watson, 1998; Bilde *et al*., 2007; Welke & Schneider, 2009) in several species. We would have expected that any direct benefit derived by mating twice, for example reception of higher amount of sperm and/or ejaculate-associated components, would be revealed in the form of increased fecundity (i.e., offspring number) for both, monogamous and polyandrous females (Arnqvist & Nilsson, 2000). Yet, only polyandrous females would experience indirect benefits in the form of enhanced fertility (i.e., higher egg hatching), by mating with males of different genetic quality and/or compatibility (Jennions & Petrie, 2000). Due to lack of visible mating plugs, it appears unlikely that plugging of the female genitalia during the first mating prevented sperm transfer during the second mating. In contrast to several other spider species which sacrifice their entire pedipalp to plug the females genitalia, *P. tepidariorum* males only can break off a very small most distal part their pedipalp which remains inside the female reproductive tract (Abalos & Baez, 1963). Whether this is enough to hinder or lower fertilization success of subsequent matings in females is however not known, and would be an interesting venue for future research.

#### Male-female behaviors at mating

The second important outcome of our study is the quantitative description of male and female behaviors at mating. Most ecological studies on *P. tepidariorum* focus primarily on prey capture and web building (Valerio, 1975; Barghusen *et al*., 1997; Hajer & Hrubá, 2007; Uma & Weiss, 2012; Brown & Houghton, 2019), leaving mating behaviors largely undescribed. Description of mating elements primarily come from two studies. The first – yet based on only 3 observations – notes that males upload sperm in their pedipalps before copulation through a lengthy process (namely, sperm induction), copulation is obtained by male approach with courtship lasting approx. 17 minutes, sperm is transferred through a total of 2 palp insertions, and mating plugs do not occur (Knoflach, 2004). In a more recent study, Ma et al. (2021) reported, in undisturbed treatment groups consisting of 20 spider pairs, a 75% mating success, 13.3% of cannibalism, and courtship (defined as the time from male vibratory performance to mating) lasting on average of 155.5 seconds. Our findings differ from the latter, as while cannibalism rates are comparable, mating rates were generally lower and time until successful copulation longer. We show that male courtship generally begins with female twanging behavior, where females pluck the web with her two front legs repeatedly, facing the direction of the male. The performance of such web plucking (twanging) behavior by females is commonly followed by the male approaching the female (Figure 3) and leads to successful mating. Indeed, females performed this behavior more intensively during trials in which males and females eventually successfully copulated (Figure 4). On the contrary, it was weakly performed during trials in which females eventually cannibalized the male. This behavior may be interpreted as a signaling behavior from females that advertise their receptivity. Females that did not have a functional web and were impaired in performing twanging were also not approached by males, indicating that vibrational communication via the web plays a crucial role in male-female reproductive communication in this species (Herberstein *et al*., 2014), in line with previous observations (Knoflach, 2004; Ma *et al*., 2022). Males responded to female signaling, by approaching the female, through jerking movements and contact movements of their legs, not always being successful in entering the mating position, as multiple copulation attempts were observed during a single trial (as discussed above). This is an interesting pattern which however differs from knowledge on other comb footed spiders, where it is suggested that males first approach the mate (Knoflach, 2004).

### Conclusions

Low mating rates as revealed for this species, suggest that high selective pressures potentially reasoned by ecological factors and mating risks for males, are at play to limit mating opportunities. This is further suggested by the lack of fitness benefits derived from multiple matings. While not affecting female reproductive output, matings may carry risks to males due to cannibalistic females. Males may therefore, be under selection to discriminate female receptiveness (Bonduriansky, 2001), most likely through multi-modal sensory channels, including chemical and vibrational web-bound communication. Lastly, this study shows that *Parasteatoda tepidariorum* is not only a good developmental model organism amenable to various established techniques to study gene expression and function, but also serves well as a behavioural, neural, and evolutionary model species, given its interesting communication and mating behavior and its short generation time and high numbers of offspring produced by one single mating.

## Acknowledgements and Funding

We thank Felix Simon Christian Quade for useful discussion on the study, Magdalena Ines Schacht for instructing on egg counting and Michelle Beyer for assistance in using R. CT was funded by the LMUexcellent Junior Research Fund

## Author contributions and competing interest statement

CT and NT designed and supervised the study. AA performed all experiments, scored video recordings and performed statistical analysis of the data under CT’s supervision. CT and NT wrote the manuscript, with AA’s contribution. All authors have read and approved the final version of the manuscript, and declare no competing interests.

## Appendices

**Table S1.** Results from the expanded models including male age (N of adulthood days) and spider origin (from wild caught, W and genome line, L), investigating a) the effects of mating treatment (polyandry and monogamy), mating trial (1 and 2) and their interaction, and relative body mass between the sexes (female-male mass), on mating success of the pair (GLMM binomial) and **b)** of mating treatment (polyandry and monogamy) and the outcome of the previous mating trial (successful, unsuccessful mating) on the likelihood of re-mating during trial 2 (LM-binomial). Significant effects are shown in bold.

|  | **a) Mating success** | **b) Re-mating success** |
| --- | --- | --- |
| **Fixed effects** | **$\beta$ (95% CI)** |  |
| (Intercept)* | -0.86<br>(-1.84 , 0.11) | -1.54<br>(-2.69 , -0.39) |
| Treatment (polyandry) | **1.68**<br>**(0.45 , 2.94)** | 0.14<br>(-0.97 , 1.28) |
| Mating trial (1,2)<sup>a</sup> | -0.59<br>(-1.66 , 0.55) |  |
| Treatment X Mating trial<sup>a</sup> | -1.23<br>(-2.78 , 0.23) |  |
| Relative Body Mass<sup>a</sup> | -2.11<br>(20.24 , 14.53) | -0.64<br>(-19.55 , 18.69) |
| Male origin (W, G) W | 0.23<br>(-0.96 , 1.46) | 0.66<br>(-0.74 , 2.06) |
| Female origin (W, G) W | 0.09<br>(-1.02 , 1.14) | 0.28<br>(-0.96 , 1.59) |
| Male age<sup>b</sup> | 0.20<br>(-0.39 , 0.83) | 0.09<br>(-0.61 , 0.77) |
| Previously mated (yes, no) |  | 0.59<br>(-0.37 , 1.58) |
| **Random Effects** | **$\sigma^2$(CI 95%)** |  |
| Female ID | 1.23<br>(0.89 , 1.62) |  |
| Male ID | 0<br>(0 , 0) |  |

\*Reference category, estimate for treatment monogamy and mean values of remaining fixed effects
<sup>a</sup> mean centered
<sup>b</sup> mean-centered and normalized with the standard-deviation

**Table S2.** Results from the expanded models including male age (N of adulthood days) and spider origin (from wild caught, W and genome line, L), investigating the effects of mating treatment (polyandry and monogamy), mating trial (1 and 2) and their interaction, relative body mass difference (female-male mass), on male and female mating behaviors.

|  | Time to female twanging | Total N of female twanging | Time to 1<sup>st</sup> male approach | Total N male approaches | Time to mating |
| --- | --- | --- | --- | --- | --- |
| **Fixed Effects** | **$\beta$ (95% CI)** |  |  |  |  |
| (Intercept)* | 4.85<br>(4.05 , 5.63) | 3.18<br>(2.57 , 3.78) | 5.41<br>(4.69 , 6.17) | 1.63<br>(0.98 , 2.26) | 415.08<br>(14.13 , 777.73) |
| Treatment (polyandry) | 0.24<br>(-0.49 , 0.94) | 0.07<br>(-0.52 , 0.62) | -0.09<br>(-0.77 , 0.59) | 0.32<br>(-0.28 , 0.89) | -12.73<br>(-306.97 , 282.36) |
| Mating trial (1,2)<sup>a</sup> | 0.07<br>(-0.02 , 0.16) | -0.05<br>(-0.16 , 0.07) | 0.11<br>(0.03 , 0.19) | -0.26<br>(-0.45 , -0.04) | 19.59<br>(-305.39 , 332.07) |
| Treatment x Mating trial<sup>a</sup> | 0.79<br>(0.67 , 0.92) | -0.47<br>(-0.70 , -0.24) | 0.89<br>(0.75 , 1.04) | -0.61<br>(-1.02 , -0.19) | -286.22<br>(-853.01 , 267.01) |
| Relative Body Mass<sup>a</sup> | -9.30<br>(-22.23 , 3.57) | 3.20<br>(-6.49 , 12.62) | 2.56<br>(-9.14 , 13.81) | 0.11<br>(-10.25 , 10.13) | -991.48<br>(-7434.23 , 5585.02) |
| Male origin (W, G) W | -0.87<br>(-2.11 , 0.39) | -0.41<br>(-1.46 , 0.63) | -0.71<br>(-1.92 , 0.51) | -0.12<br>(-1.24 , 0.97) | 195.39<br>(-440.84 , 864.15) |
| Female origin (W, G) W | -1.04<br>(-2.06 , 0.02) | 0.49<br>(-1.31 , 0.30) | -0.68<br>(-1.63 , 0.32) | -0.37<br>(-1.23 , 0.48) | 15.75<br>(-472.93 , 542.12) |
| Male age<sup>b</sup> | -0.21<br>(-0.70 , 0.30) | -0.03<br>(-0.45 , 0.39) | -0.17<br>(-0.65 , 0.31) | 0.03<br>(-0.41 , 0.47) | -286.22<br>(-174.47 , 353.55) |
| **Random Effects** | **$\sigma^2$(CI 95%)** |  |  |  |  |
| **Female ID** | 0.89<br>(0.61 , 1.24) | 0.27<br>(0.18 , 0.37) | 0.48<br>(0.31 , 0.69) | 0.27<br>(0.18 , 0.37) | 0<br>(0 , 0) |
| **Male ID** | 0.95<br>(0.71 , 1.26) | 0.89<br>(0.65 , 1.16) | 1.05<br>(0.75 , 1.43) | 0.80<br>(0.58 , 1.08) | 17910.29<br>(8007.63 , 35116.29) |

\*Reference category, estimate for treatment monogamy and mean values of remaining fixed effects
<sup>a</sup> mean centered
<sup>b</sup>mean-centered and normalized with the standard-deviation

**Table S3.** Results from the GLMs investigating the effects of mating treatment (mated once, twice M, twice P) and female body mass on fitness.

|  | Total number of eggs laid per cocoon |  | Proportion of viable eggs per cocoon |  |
| --- | --- | --- | --- | --- |
| | | $\beta$ (95% CI) | | |
| **(Intercept)\*** | 149.91<br>(105.45 , 193.59) | 152.18<br>(107.26 , 197.730) | 0.55<br>(0.39 , 0.69) | 0.53<br>(0.38 , 0.68) |
| **Mated twice monogamy** | -16.58<br>(-53.55 , 19.26) | -17.19<br>(-54.07 , 18.79) | -0.06<br>(-0.21 , 0.07) | -0.06<br>(-0.19 , 0.08) |
| **Mated twice polyandry** | 13.17<br>(-14.88 , 43.02) | 13.82<br>(-16.03 , 43.67) | -0.10<br>(-0.21 , 0.02) | -0.11<br>(-0.22 , 0.008) |
| **Female Body Mass** | 331.38<br>(-60.62 , 747.57) | 337.78<br>(-110.97 , 761.99) | 1.37<br>(-0.17 , 3.00) | 1.32<br>(-0.40 , 2.95) |
| **N cocoon (1-4)** | -2.19<br>(-10.79 , 6.34) | -2.26<br>(-10.29 , 6.02) | 0.02<br>(-0.004 , 0.04) | 0.02<br>(-0.002 , 0.04) |
| **Female origin (W, G) W** |  | -6.76<br>(-29.02 , 15.19) |  | 0.06<br>(-0.03 , 0.14) |
| **Random Effects** | | $\sigma^2$(CI 95%) | | |
| **Female ID** | 373.72<br>(224.89 , 563.97) | 391.48<br>(236.44 , 599.64) | 0.01<br>(0.009 , 0.02) | 0.01<br>(0.008 , 0.02) |

\*Reference category, estimate for treatment monogamy and mean values of remaining fixed effects

## Notes

### Competing Interest Statement

The authors have declared no competing interest.

